# Anaerobic conditions increase plasmid transfer rates across *Escherichia coli* strains

**DOI:** 10.64898/2026.06.25.734511

**Authors:** Ingrid Cárdenas-Rey, Sally Felle, Michael S. M. Brouwer, Kees T. Veldman, J. Arjan G. M. de Visser

## Abstract

Bacterial conjugation is the primary mechanism by which antibiotic resistance genes spread in microbial populations, yet our understanding of this process has been largely based on experiments conducted under aerobic conditions. This creates a fundamental disconnect: environments that are considered hotspots for gene exchange (e.g., the gut, abscesses, chronic wounds, and wastewater systems) are predominantly anaerobic. In this study, we investigate whether oxygen availability influences the transfer rate of a set of common ESBL-IncI1-α and *qnrS1*-IncF plasmids in commensal *Escherichia coli* strains from chickens. We found that oxygen availability significantly shapes conjugation dynamics in a recipient strain-specific manner, with anaerobic conditions promoting higher ESBL-IncI1-α plasmid transfer rates to commensal *E. coli* recipients. Conjugation rates of the ESBL-IncI1-α plasmids to a laboratory strain of *E. coli* were several orders of magnitude higher and independent of oxygen level, while two *qnrS1*-IncF plasmids showed higher anaerobic rates. Our study reveals critical “oxygen blind spots” in conventional conjugation assays and suggests that conventional aerobic conjugation assays underestimate plasmid transfer rates in natural environments such as the chicken caeca. These findings highlight the importance of aligning experimental conditions with the physiological and ecological environments in which gene exchange naturally occurs. Tailoring these variables is essential for generating results that accurately reflect, predict, and potentially intervene in the horizontal spread of antimicrobial resistance.

## INTRODUCTION

Bacterial conjugation is a major driver of horizontal gene transfer (HGT), facilitating the spread and maintenance of antimicrobial resistance (AMR) genes and other adaptive traits across diverse bacterial populations (Devanga Ragupathi et al., 2019; Shen et al., 2022; T. Wang & You, 2020; X. Wang et al., 2023). HGT is successfully enabled by conjugative plasmids and other mobile genetic elements, which encode the machinery for their own transfer and are frequently implicated in the dissemination of clinically relevant resistance determinants (Alonso-del Valle et al., 2023; Norman et al., 2009). Despite the central role of bacterial conjugation in the spread of antimicrobial resistance, relatively few studies have systematically examined the influence of experimental conditions on plasmid transfer rates (Alderliesten et al., 2020; Huang et al., 2022; Johnsen & Kroer, 2007; Neil et al., 2020a; Pallares-Vega et al., 2021). Most experimental *in vitro* conjugation designs rely on aerobic static cultures incubated at 37 °C, which, while convenient, often do not reflect the physiological environments where gene exchange frequently occurs (Dahl et al., 2007; Ding et al., 2025; Neil et al., 2020a; Ott et al., 2020; Ott & Mellata, 2024; Stecher et al., 2012; Zeng et al., 2026).

Conjugation experiments vary widely in protocol, including the use of filter versus liquid matings and different enumeration methods, which makes cross-study comparisons difficult (He et al., 2024). Reported outcome metrics also differ: transconjugant frequencies and transconjugant-to-donor ratios are commonly used, yet both depend strongly on the donor and recipient cell densities, because these determine the number of mating opportunities. Despite this, only a few studies account for changes in mating opportunities during the assay, even though donor and recipient densities typically shift over time (Benz & Hall, 2023; Duxbury et al., 2021). As a result, widely used metrics often capture the density-driven encounter process between donors and recipients rather than the underlying plasmid transfer rate (Huisman et al., 2022).

Anaerobic conditions, characteristic of niches such as the large intestine, abscesses, chronic wounds, and wastewater systems, which serve as natural reservoirs for gene exchange (Aviv et al., 2016; Licht et al., 1999; Neil et al., 2020; Smyth et al., 2025; Sroithongkham et al., 2025; Zubair et al., 2012), are particularly underrepresented in *in vitro* conjugation assays. A limited number of studies have reported lower conjugation frequencies under anaerobic conditions (Ochi et al., 2021; Piscon et al., 2023). However, because they do not incorporate changes in donor and recipient densities over time, their estimates primarily capture density-dependent encounter dynamics rather than the biological rate of plasmid transfer. These gaps raise concerns that prevailing conjugation estimates may misrepresent *in vivo* dynamics, thereby leading to inaccurate assumptions within comparative risk assessment of plasmid-mediated AMR-gene transfer. Addressing this gap is critical for generating data that better approximates natural transfer conditions and contextualises laboratory findings within clinically relevant settings.

In this study, we use a factorial experimental design with commensal *E. coli* donors and recipients originating from the caeca of chickens. We compare transfer rates of three ESBL-IncI1-α plasmids at 41°C (the chicken’s body temperature) to three recipient strains under both anaerobic (caeca’s natural oxygen levels) and aerobic (atmospheric) conditions. Our findings suggest that oxygen availability modulates conjugation rates in a recipient strain-specific manner, with generally higher conjugation rates under anaerobic conditions. Additionally, using commensal *E. coli* donor strains carrying either ESBL-IncI1-α or *qnrS1*-IncF plasmids together with a universal K12-*yfp* laboratory recipient, we demonstrate that the two plasmid types exhibit distinct oxygen-dependent transfer behaviours. ESBL-IncI1-α plasmids show oxygen-independent conjugation, maintaining consistently high transfer rates in both aerobic and anaerobic conditions, whereas *qnrS1*-IncF plasmids achieve substantially higher conjugation rates under anaerobic conditions. Our results highlight the importance of considering the specific environmental context (e.g., oxygen availability) and strain and plasmid traits when designing and performing bacterial conjugation assays to accurately assess the dynamics of plasmid-mediated AMR dissemination.

## RESULTS

### Oxygen availability and recipient strain shape conjugation outcomes

To assess whether oxygen availability influences ESBL-IncI1-α plasmid transfer, we used a full factorial design, in which three chicken-commensal *E. coli* ESBL-carrying donors (D1-D3) were each paired with three ciprofloxacin-resistant chicken-commensal recipients (R1-R3), yielding nine donor–recipient combinations (Table 1). Liquid matings were performed under aerobic and anaerobic conditions, with three biologically independent replicates conducted on separate days using freshly prepared bacterial cultures. Plasmid transfer rates were estimated using the Approximate Extended Simonsen Model (Huisman et al., 2022), which incorporates donor and recipient population dynamics to infer the underlying transfer rate rather than density-dependent outcome measures.

**Table 1.** Plasmids and resistance genes of chicken-commensal donor and recipient strains used in this study.

| Isolate ID | Type<sup>a</sup> | Resistance genes | Plasmid | Reference<sup>b</sup> |
| --- | --- | --- | --- | --- |
| 33-68 | D1 | *bla*<sub>CTX-M-1</sub> | pESBL33-68: IncI1-α | (Brouwer et al., 2023) and this study |
| 34-08 | D2 | *bla*<sub>CTX-M-1</sub> | pESBL34-08: IncI1-α | (Brouwer et al., 2023) and this study |
| 34-22 | D3 | *bla*<sub>CTX-M-1</sub> | pESBL34-22: IncI1-α | (Brouwer et al., 2023) and this study |
| 271-15 | D4 | *qnrS1* | pQNR271-15:<br>IncFIB(AP001918)/IncFIC(FII) | This study |
| 274-77 | D5 | *qnrS1* | pQNR274-77:<br>IncFIB(AP001918)/IncFIC(FII) | This study |
| 278-15 | D6 | *qnrS1* | pQNR278-15:<br>IncFIA/IncFIB(AP001918)/IncFIC(FII) | This study |
| 208-10 | R1 | QRDR<sup>c</sup> | - | This study |
| 259-14 | R2 | QRDR | - | This study |
| 270-23 | R3 | QRDR | - | This study |
| K12-*yfp* | R4 | *catA1* | - | (Elowitz et al., 2002) and this study |

<sup>a</sup>D1-D6; *E. coli* commensal donor strains from chickens. R1-R3, *E. coli* commensal recipient strains from chickens.
<sup>b</sup>D1- D3: DNA short-read sequencing was performed by Brouwer et al. (2023), and long-read sequencing was performed in this study. D4-D6 and R1-R3: DNA sequencing was performed in this study. K12-*yfp* was created by Elowitz et al. (2002), and DNA sequencing was performed in this study.
QRDR; Quinolone Resistance-Determining Region recipients (chromosomal mutations in *gyrA* and *parC*)

Plasmid transfer rates across all nine donor–recipient pairs spanned several orders of magnitude both under aerobic and anaerobic environments. However, most donor–recipient pairs showed consistently higher transfer rates in the anaerobic environment. Notably, all pairs containing R3 displayed the highest transfer rates overall (Fig. 1A).

**Figure 1.**
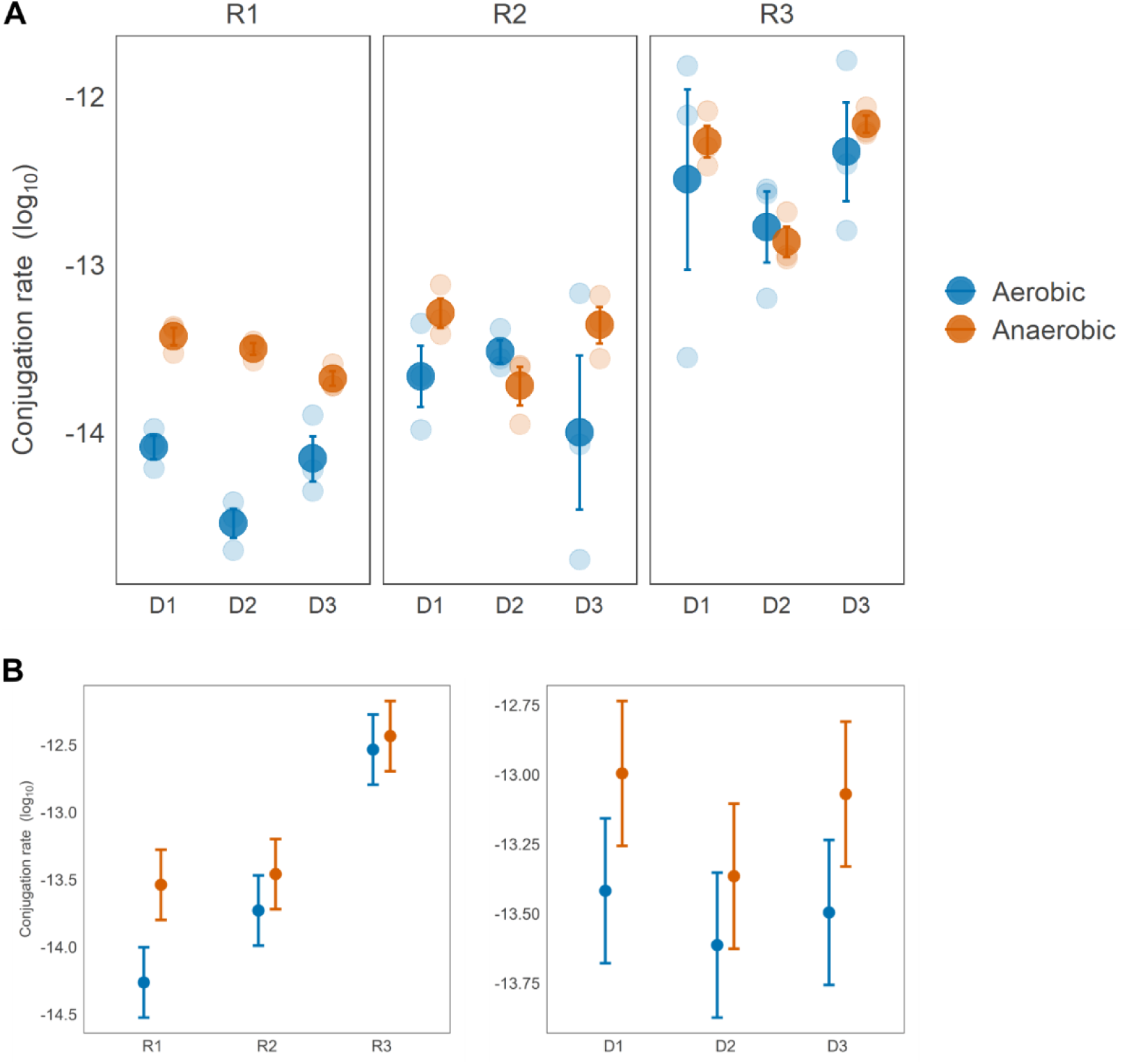
Conjugation rates of ESBL–IncI1-α plasmids are shaped by recipient strain and oxygen availability. **A.** Plasmid transfer rates [mL (CFU h)^−1^] for nine ESBL donor–QRDR recipient pairs under aerobic (blue) and anaerobic (orange) conditions. Light dots represent independent biological replicates performed on separate experimental days (*n* = 3); dark dots and error bars indicate mean ± SE. Conjugation rates were calculated using the Approximate Extended Simonsen Model. **B.** Estimated marginal means of conjugation rates [mL (CFU h)−1] for recipients averaged across all donors (left), and donors averaged across all recipients (right) under aerobic and anaerobic conditions. Dots denote means; error bars indicate 95% confidence intervals.

We used a linear mixed-effects model to quantify the effects of oxygen availability (anaerobic vs. aerobic environment), donor strain, and recipient strain on conjugation rates. The model included fixed effects for all three factors (donor, recipient and environment) and their two-way and three-way interactions, with independent biological replicates as a random intercept to control for day-to-day biological variation. Type III ANOVA applied to the model revealed strong and structured variation in conjugation rates across factors. The environment had a significant main effect, with higher conjugation rates under anaerobic conditions. Recipient strain was the strongest predictor, explaining 68.67% of the variance in conjugation rates, whereas donor strain showed only a marginal effect. Although recipients accounted for the largest share of variance in conjugation rates, this effect was environment-dependent, as indicated by the significant recipient–environment interaction (Table 2).

**Table 2.** Effects of environment (aerobic vs anaerobic), donor, and recipient strain on ESBL-Inc1α conjugation rates.

| Term<sup>a</sup> | SS<sup>b</sup> | MS<sup>c</sup> | DF<sub>Num/Den</sub><sup>d</sup> | F<sup>e</sup> | P | Explained variance (%)<sup>f</sup> |
| --- | --- | --- | --- | --- | --- | --- |
| Environment | 1.80 | 1.80 | 1,34 | 15.08 | **4.5e-04** | 6.19 |
| Donor | 0.77 | 0.38 | 2, 34 | 3.22 | 0.052 | 2.64 |
| Recipient | 19.98 | 9.99 | 2, 34 | 83.62 | **7.4e-14** | 68.67 |
| Environment x Donor | 0.09 | 0.05 | 2, 34 | 0.39 | 0.679 | 0.32 |
| Environment x Recipient | 0.94 | 0.47 | 2, 34 | 3.93 | **0.029** | 3.22 |
| Donor x Recipient | 0.66 | 0.16 | 4, 34 | 1.38 | 0.262 | 2.27 |
| Environment x Donor x Recipient | 0.79 | 0.20 | 4, 34 | 1.66 | 0.182 | 0.18 |
| Residuals |  |  |  |  |  | 13.96 |

<sup>a</sup>Type III ANOVA results from a linear mixed-effects model fitted to log<sub>10</sub>-transformed conjugation rates from the full factorial experiment (3 ESBL donors × 3 QRDR recipients × 2 environments). Fixed effects included environment (aerobic vs. anaerobic), donor strain, recipient strain, and all two-way and three-way interactions. A random intercept for experimental day was included to account for temporal variation across independent replicates. Significant *p*-values (*p* < 0.05) are shown in bold. <sup>b</sup>SS, sum of squares; <sup>c</sup>MS, mean squares; <sup>d</sup>DF<sub>Num, Den</sub>, numerator and denominator degrees of freedom calculated using Satterthwaite's approximation; <sup>e</sup>F, F-statistic; <sup>f</sup>Explained variance (%), proportion of the total sum of squares for each fixed term and residual.

The random intercept for independent biological replicates captured a small amount of between-day variability, indicating that baseline conjugation rates differed slightly across days relative to the fixed effects and did not drive the observed environment- and recipient-specific patterns (Table S1).

Estimated marginal means further elucidated the nature of the environment–recipient interaction. Averaged over donors, R1 exhibited the lowest conjugation rate in the aerobic environment but showed an approximate five-fold increase in the anaerobic environment. R2 showed a modest, non-significant increase from aerobic to anaerobic environment, whereas R3 maintained consistently high conjugation rates without sensitivity to oxygen availability (Fig. 1B). Pairwise Tukey-adjusted contrasts for recipient-environment combinations showed that under aerobic conditions, R1 had significantly lower conjugation rates than both R2 (*p* = 0.007) and R3 (*p* <0.001). However, anaerobically, R1 and R2 no longer differed, but both exhibited significantly lower conjugation rates than R3 (*p* <0.001). Together, these results suggest that oxygen availability modulates conjugation rates in a recipient-specific manner, with R1 being strongly oxygen-sensitive, R2 moderately affected, and R3 largely oxygen-independent (Fig. S1).

### Oxygen availability drives divergent plasmid-dependent shifts in conjugation rates

To further explore the effect of oxygen availability on plasmid transfer, we performed additional conjugation assays using the same experimental conditions described above, this time employing a well-characterised *E. coli* K-12 laboratory recipient strain. This strain was chosen to avoid interference from resident plasmids and minimise variability arising from recipient-specific defence mechanisms (see below). The K-12 recipient was mated with six commensal *E. coli* donors, including the three strains carrying ESBL-IncI1-α plasmids involved in the experiments described above (D1-D3) and three strains carrying *qnrS1-*IncF plasmids (D4-D6).

Across both environments, conjugation rates varied over several orders of magnitude. The three ESBL-IncI1-α donors transferred at rates that were at least two to five orders of magnitude higher than in the experiments involving commensal strains as recipients. In addition, oxygen availability hardly affected transfer rates (only D2 showed slightly higher anaerobic rates), while *qnrS1*-IncF donors (D4 and D6) showed a pronounced increase in transfer rates specifically under anaerobic conditions (Fig. 2). Donor D5 was excluded from statistical analyses as it did not yield transconjugant cells.

**Figure 2.**
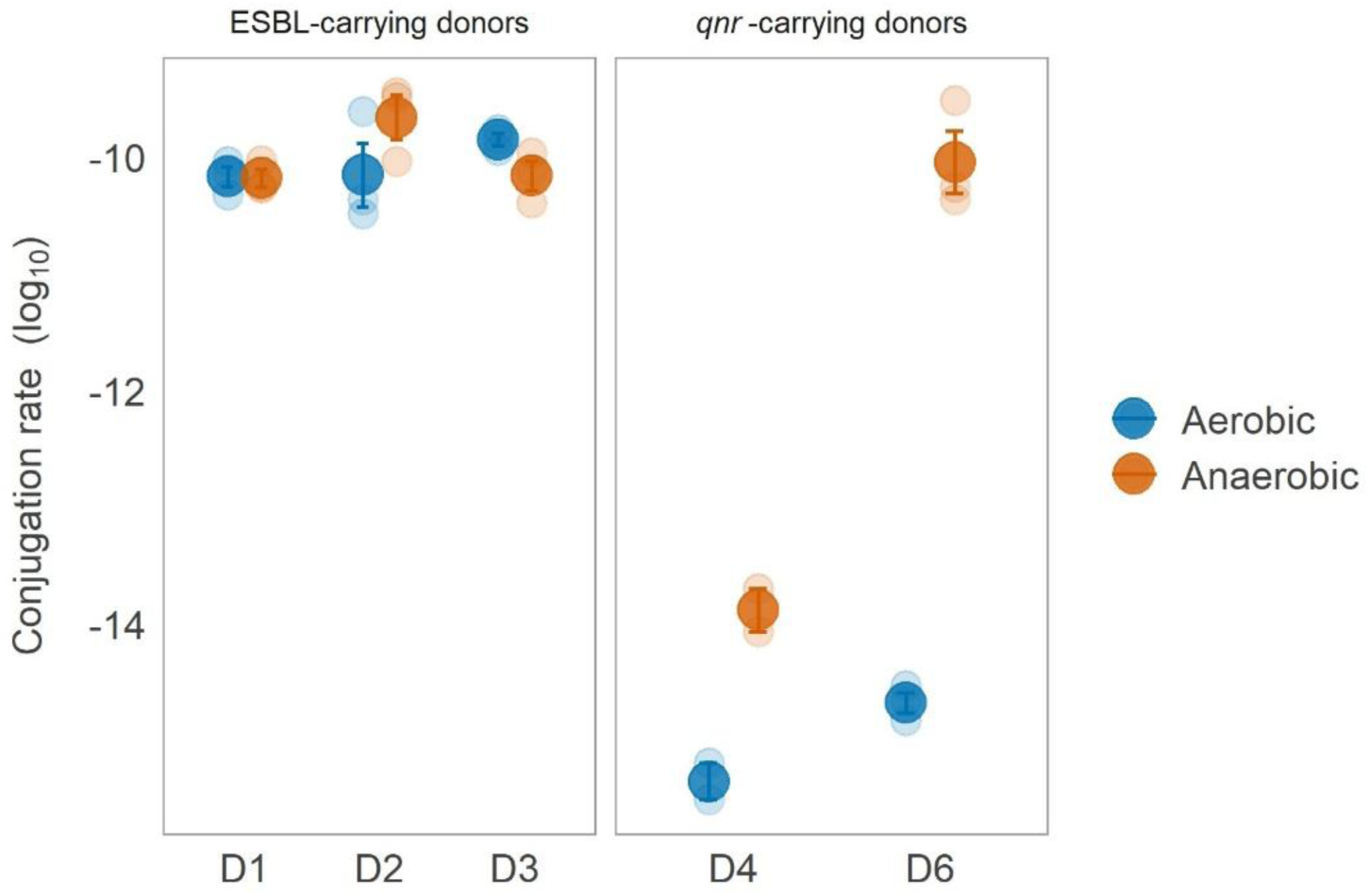
Oxygen-dependent conjugation rates of ESBL-IncI1-α and *qnrS1*-IncF plasmids to recipient *E. coli* K-12. Plasmid transfer rates [mL (CFU h)^−1^] for each donor–recipient pair under aerobic and anaerobic conditions. Light dots represent independent biological replicates performed on separate experimental days (n = 3 per pair); dark dots and error bars indicate mean ± SE. Conjugation rates were calculated using the Approximate Extended Simonsen Model.

We then quantified the effects of oxygen availability and plasmid type on conjugation rates using a linear mixed-effects model that included environment (aerobic vs. anaerobic) and plasmid type as fixed effects, their interaction, and a random intercept for donor strain to account for donor-specific biological variation. A significant plasmid type–specific response to oxygen availability was observed (Table 3). *qnrS1*-IncF plasmids showed markedly higher transfer rates under anaerobic conditions (Fig. S2), whereas ESBL-IncI1-α plasmids exhibited high conjugation rates across both environments. The random-effects estimates showed minimal donor-level variation, indicating that donors did not substantially influence baseline conjugation rates (Table S2).

**Table 3.** Effects of oxygen availability and plasmid type on conjugation rates.

| Term<sup>a</sup> | SS<sup>b</sup> | MS<sup>c</sup> | DF<sub>Num, Den</sub><sup>d</sup> | F<sup>e</sup> | P | Explained variance (%)<sup>f</sup> |
| --- | --- | --- | --- | --- | --- | --- |
| Environment | 20.04 | 20.04 | 1, 20.00 | 206.093 | **9.4e-10** | 24.86 |
| Plasmid type | 72.35 | 36.17 | 2, 2.55 | 372.032 | **3.7e-07** | 16.28 |
| Environment x Plasmid type | 23.57 | 11.78 | 2, 20.00 | 121.191 | **5.8e-11** | 55.87 |
| Residuals |  |  |  |  |  | 2.98 |

<sup>a</sup>Type III ANOVA results from a linear mixed-effects model fitted to log<sub>10</sub>-transformed conjugation rates with environment (aerobic vs anaerobic) and plasmid type as fixed effects, their two-way interactions, and donor strain as a random intercept to control for donor biological variation. Significant *p*-values (*p* < 0.05) are shown in bold. <sup>b</sup>SS, sum of squares; <sup>c</sup>MS, mean squares; <sup>d</sup>DF<sub>Num, Den</sub>, numerator and denominator degrees of freedom calculated using Satterthwaite's approximation; <sup>e</sup>F, F-statistic; <sup>f</sup>Explained variance (%), proportion of the total sum of squares for each fixed term and residual.

An additional linear mixed-effects model was fitted with environment and donor as fixed effects, their two-way interactions, and independent biological replicates as a random intercept to account for day-to-day biological variation (Table S3). In line with the previous model, *qnrS1*-IncF plasmid-carrying donors (D4 and D6) showed marked increases in conjugation rates anaerobically compared to their aerobic baselines, whereas among ESBL-IncI1-α donors, only D2 exhibited a modest anaerobic increase (Table S4).

Donors D4 and D6, both carrying *qnrS1*-IncF plasmids, showed substantially lower conjugation rates than the ESBL-IncI1-α donors D1, D2, and D3 (all *p* < 0.001) under aerobic conditions (Fig. S3). For example, D4, which carried an IncFIB/FIC(FII) plasmid, transferred at > 10⁵-fold lower rates than D1. Under anaerobic conditions, however, this gap narrowed markedly. Donor D6, which carried a multi-replicon IncFIA/FIB/FIC(FII) plasmid, reached conjugation levels comparable to all ESBL-IncI1-α donors and significantly exceeded D4 (*p* < 0.001), consistent with the strong plasmid type × environment interaction (Fig. S3).

### Diverse phylogenetic backgrounds and plasmid profiles among donor and recipient strains

All donor and recipient strains (n=10) used in this study underwent short-read (Illumina) sequencing for genome characterisation and resistance gene profiling. Long-read sequencing (Oxford Nanopore Technologies) was additionally performed on the six donor isolates to enable complete characterisation of the plasmids of interest. The isolates represented diverse phylogenetic backgrounds and sequence types (STs; see Fig. 3). Most belonged to phylogroup B1, whereas the K12-*yfp* recipient was assigned to phylogroup A and D6 to phylogroup F. Five multi-locus sequence types (MLSTs) were identified, with ST162 being the most prevalent (n=4). Notably, nearly all donor-recipient pairs belonged to different STs, except for one pair, donor D2 and recipient R1, which were both ST155 (Fig. 3).

**Figure 3.**
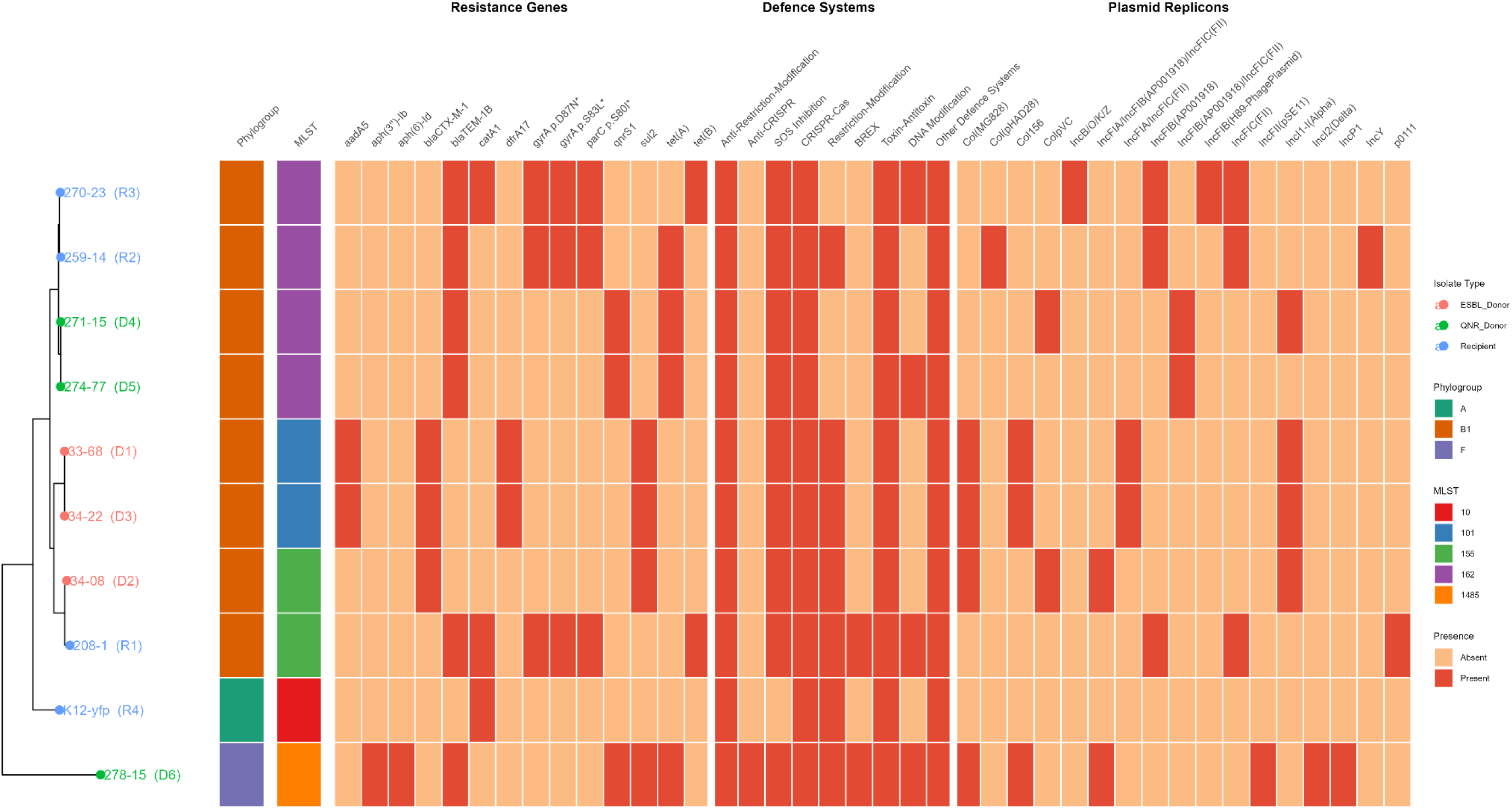
Phylogenetic relationships and molecular characteristics of donor and recipient isolates. (Left) Maximum likelihood phylogenetic tree based on core genome alignment. The tree tips are colour-coded to indicate ESBL donors (red), *qnrS1* donors (green) and recipients (blue). (Right) Heatmap showing the distribution of molecular features, including phylogroup, multilocus sequence type (MLST), antimicrobial resistance genes, defence systems relevant to plasmid conjugation, and plasmid replicons. Presence/absence is indicated by dark coral/light coral, respectively. Asterisks (*) denote chromosomal point mutations conferring antibiotic resistance.

Resistance gene profiles varied among the isolates (Fig. 3). The three commensal *E. coli* recipients R1, R2, and R3 each carried identical mutations in the quinolone resistance-determining region (QRDR), conferring high-level fluoroquinolone resistance. The K12-*yfp* recipient carried only the *catA1* gene, conferring chloramphenicol resistance, whereas all other isolates harboured multiple resistance determinants. As expected, resistance genes of interest segregated by plasmid type. The three ESBL-IncI1-α donors D1, D2, and D3, exclusively carried *bla*_CTX-M-1_, while the three *qnrS1*-IncF donors D4, D5, and D6, exclusively carried *qnrS1*.

Defence and anti-defence system profiling revealed heterogeneous distributions across isolates (Fig. 3, detailed gene list in Table S5). All isolates carried anti-restriction modification, CRISPR-Cas and toxin-antitoxin systems. Only *qnrS1*-IncF D6 carried an anti-CRISPR system, though this was not located on the plasmid of interest (pQNR278-15). Recipient R1 additionally carried a truncated Bacteriophage Exclusion (BREX) system.

PlasmidFinder analysis identified diverse incompatibility groups across the isolates (Fig. 3). K12-*yfp* carried no detectable plasmid replicons, while most other isolates harboured multiple replicons. The three ESBL donors (D1-D3) shared a common IncI1-α replicon carrying the *bla*_CTX-M-1_ gene, with two donors, D1 and D3, exhibiting identical replicon profiles. Among the *qnrS1*-IncF donors, D4 and D5 carried IncFIB(AP001918)/IncFIC(FII) replicons, while D6 carried IncFIA/IncFIB(AP001918)/IncFIC(FII). These IncF plasmids corresponded to the *qnrS1*-carrying plasmids of interest. All recipients, except for K12-*yfp*, carried multiple plasmid replicons (Fig 3).

Phylogenetic analysis revealed varied relationships among donor-recipient pairs (Fig. 3). Recipient R1 clustered with ESBL-IncI1-α donor D2, while the remaining two IncI1-α donors D1 and D3, formed a separate cluster. Recipients R3 and R2 clustered together. K12-*yfp* recipient formed its own distinct clade. Among the *qnrS1*-IncF donors, D4 and D5 clustered together, while D6 was phylogenetically distinct.

Collectively, these genomic features provide context for interpreting the conjugation results. The absence of resident plasmid replicons in K12-*yfp* is consistent with the higher conjugation rates observed for this recipient compared with the commensal strains (Fig. 1 and Fig. 2). Among the commensal recipients, resistance gene profiles and replicon content were broadly similar, and no single genomic feature clearly explained the strong recipient-dependent variation in conjugation rates. Phylogenetic relatedness also did not predict transfer rates; for example, the shared ST155 background of R1 and D2 was not associated with enhanced conjugation in either environment. The presence of certain defence systems (including the BREX system in R1) and resident plasmid replicons across recipient isolates remains a plausible contributor to this variation.

### Donor plasmids are structurally conserved within incompatibility groups

Complete sequences of the six donor plasmids were obtained and analysed using progressiveMauve alignment to assess structural organisation and conjugation-relevant features. The three ESBL-IncI1-α plasmids showed extensive synteny and a common structural organisation (Fig. 4A). All three carried *bla*_CTX-M-1_ and *sul2*, with pESBL33-68 and pESBL34-22 containing additional resistance genes (Table 4). Each encoded a complete conjugative transfer region, including the characteristic IncI1 dual pilus system (*tra*/*trb* and *pilJ-V*), alongside counter-defence-associated genes such as SOS inhibition (*psiA* and *psiB*), and anti-restriction modification genes (*ardA* and *ardB*).

**Figure 4.**
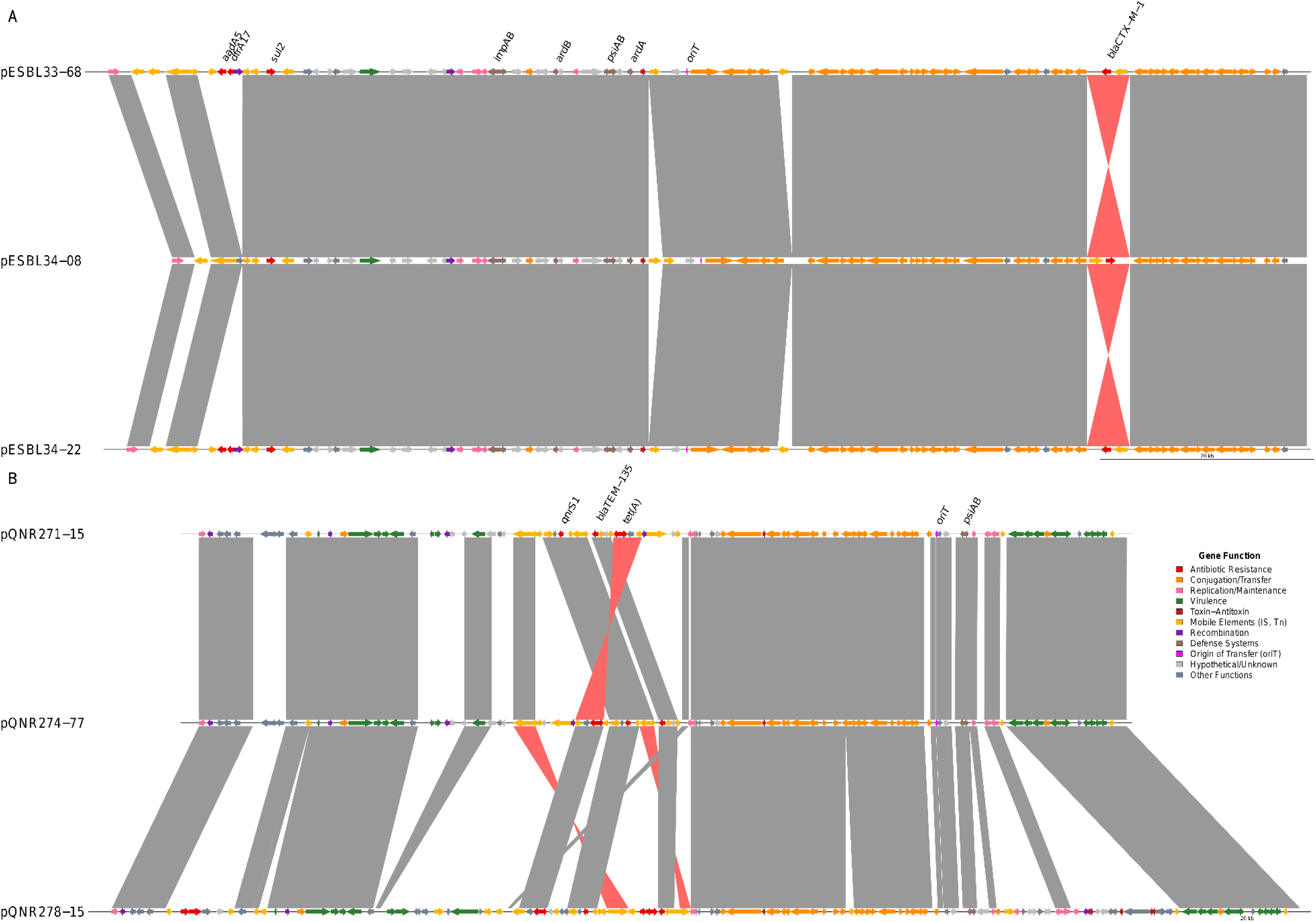
Comparative synteny of donor plasmids. Linear comparative map of (A) three ESBL-carrying plasmids (pESBL33-68, pESBL34-08, pESBL34-22) and (B) three *qnrS1*-carrying plasmids (pQNR271-15, pQNR274-77, pQNR278-15). Genes are represented as arrows, coloured by function: antibiotic resistance (red), conjugation and transfer (orange), replication/maintenance (pink), virulence factors (green), toxin-antitoxin systems (dark red), mobile elements including insertion sequences (IS) and transposons (Tn) (yellow), recombination (purple), defence systems (brown), origin of transfer (oriT) (magenta), hypothetical/unknown (dark grey), and other functions (light grey). Selected genes from key functional categories are labelled, including antibiotic resistance genes, defence system components and oriT. Grey shaded regions between plasmids indicate homologous sequence blocks in the same orientation, while red shaded regions denote inversions. Scale bars represent 20kb.

**Table 4.**
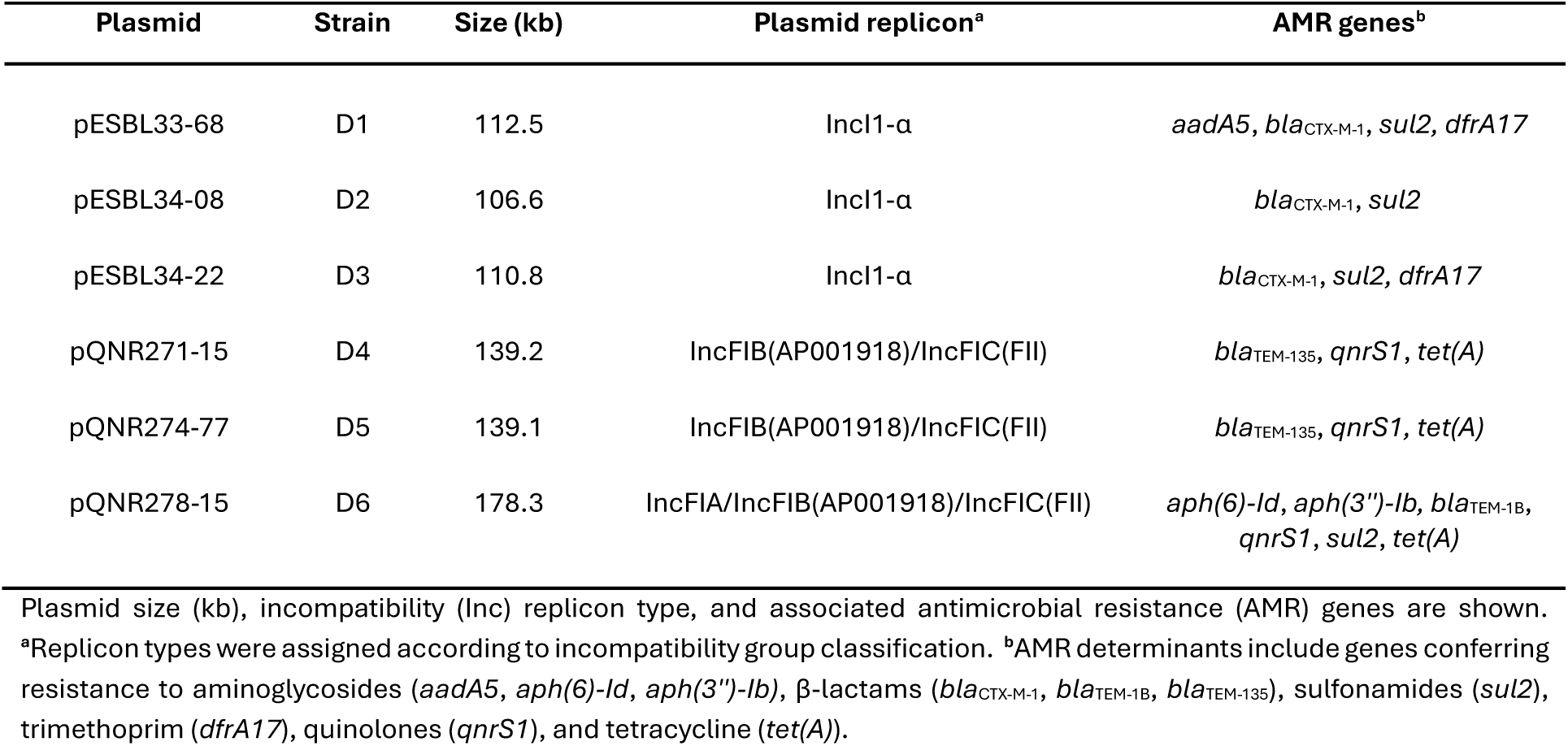
Genetic features of plasmids used in this study.

The three *qnrS1*-IncF plasmids showed high structural conservation, despite size variation (Fig 4B). The plasmid replicon for two of the plasmids (pQNR271-15 and pQNR274-77) was IncFIB(AP001918)/IncFIC(FII), while for pQNR278-15 it was IncFIA/IncFIB(AP001918)/IncFIC(FII) (Table 4). All three plasmids encoded complete conjugative transfer machinery, including a type IV secretion system (*tra*/*trb* genes) and SOS inhibition genes (*psiA* and *psiB*) within the leading transfer region (Fig. 4B). The larger size of pQNR278-15 relative to the other two IncF plasmids reflected its expanded resistance gene content and additional IncFIA replicon.

Overall, plasmids within each incompatibility group shared a common evolutionary origin, with IncF plasmids showing greater structural variability than IncI1α plasmids. The contrasting oxygen responses observed in the conjugation assays with recipient K12, where IncF plasmids showed enhanced transfer under anaerobic conditions while IncI1α plasmids did not, may partly reflect differences in the regulation of transfer genes between these plasmid types (see discussion). Among the three ESBL-IncI1α donors, only D2 exhibited a modest increase in conjugation rates under anaerobic conditions, and it was also the only ESBL donor carrying an additional IncFIA/FIB/FIC(FII) plasmid.

## DISCUSSION

Bacterial conjugation assays are a cornerstone of microbial genetics and are widely used to characterise the mobility of plasmids and their role in the dissemination of AMR across diverse bacteria and environments (Grote, 2008; Helinski, 2022; Kosterlitz & Huisman, 2023). Classical conjugation assays are typically conducted under convenient laboratory conditions, aerobic atmosphere at 37 °C, to ensure reproducibility and optimal growth conditions. However, these “ideal” parameters often fail to reflect the physiological stresses and ecological nuances of clinical or natural habitats, such as the hypoxic, nutrient-limited environment of the large intestine (Benz et al., 2021; Gumpert et al., 2017; López et al., 2025).

To investigate whether oxygen availability affects conjugation, we used a fully factorial conjugation design with commensal *E. coli* isolates from the caeca of chickens. We compared ESBL-IncI1-α plasmid transfer rates under anaerobic (mimicking the caecal environment) and aerobic (atmospheric) conditions. Anaerobic conditions significantly enhanced plasmid transfer, but this effect was strongly recipient dependent. Using donors carrying ESBL-IncI1-α or *qnrS1*-IncF plasmids with a universal K12-*yfp* recipient, we further identified divergent responses between plasmid types. Together, these findings reveal that both recipient strain and plasmid type critically shape how oxygen availability modulates conjugation rates. Since oxygen availability has wide-ranging effects on *E. coli* physiology, the variation we observed may result from effects during multiple stages of the conjugation process, including mating pair formation, DNA transfer, and plasmid stability, rather than any single mechanism (Douarche et al., 2009; Salmon et al., 2005).

The anaerobic enhancement of conjugation was not uniform across recipient strains, indicating that host factors also shape plasmid acquisition. Genomic characterisation of the isolates revealed that they all carry toxin-antitoxin systems, which influence plasmid maintenance and stability, and CRISPR-Cas systems that can restrict horizontal gene transfer by targeting foreign DNA (Garneau et al., 2010; Jurėnas et al., 2022). Beyond these shared systems, recipients differ in the presence of BREX and RM systems. BREX has recently been shown to confer anti-plasmid defence activity (Jiang et al., 2024). These differences and the presence of additional plasmid replicons in recipient strains may help to explain the observed conjugation phenotypes.

Recipient R3 consistently showed the highest conjugation rates regardless of donor or oxygen environment, a pattern that might be explained by the absence of RM systems in this strain. RM systems cleave incoming foreign DNA and are well-established barriers to conjugation (Shaw et al., 2023; Siedentop et al., 2024). All ESBL-IncI1-α plasmids carried anti-RM systems, which likely contributed to their successful transfer across the other recipients under both oxygen conditions (Dimitriu et al., 2024). Although the donor effect was small, its potential role in shaping the oxygen response cannot be excluded.

Previous studies have highlighted the influence of strain-specific factors on ESBL-plasmid spread (Benz et al., 2021). We extend this knowledge by demonstrating that experimental conditions, specifically oxygen availability, play an important role in shaping conjugation rates of ESBL-Incl1-α plasmids in a recipient-dependent manner. Past work has reported higher conjugation coefficients of ESBL-IncI plasmids in low-oxygen environments (e.g., the chicken caeca) compared with aerobic *in vitro* settings (Fischer et al., 2019; Licht et al., 1999). Neil et al. (2020) similarly observed ∼10-fold higher transfer of IncI2 plasmids under low-oxygen conditions in a murine model. However, oxygen availability has often been overlooked as a potential driver among the many abiotic and host-plasmid factors influencing plasmid transfer rates.

Beyond recipient-dependent variation, we identified a striking difference in oxygen responsiveness between the two plasmid types examined. ESBL-IncI1-α plasmids showed largely oxygen-independent transfer with the laboratory strain, with consistently high conjugation rates under both aerobic and anaerobic conditions. In contrast, *qnrS1*-IncF plasmids showed markedly elevated conjugation rates under anaerobic conditions. This variation possibly reflects the different transfer regulatory architectures of the two plasmid types.

Transfer gene expression for IncI1-α is mostly driven by plasmid-encoded regulators, including TraB and TraC (Foley et al., 2021; Komano et al., 2000), while IncF plasmids rely on a multi-layered transcriptional cascade. This requires the cooperative binding of two activators, plasmid-encoded *traJ* and the host response regulator ArcA, to achieve high transcriptional output (Bischof et al., 2020; Lu et al., 2018; Strohmaier et al., 1998). ArcA is the response regulator of the ArcAB two-component system, where the sensor kinase phosphorylates ArcA under anaerobic conditions, increasing its DNA-binding affinity by up to 10-fold (Malpica et al., 2006; Strohmaier et al., 1998). Serna et al. (2010) demonstrated that ArcAB increases IncF transfer under microaerobic conditions through activation of the *tra* operon. Recently, it has also been shown that ArcA and TraJ bind cooperatively to promote a high level of expression of the *tra* operon (Lu et al., 2018). The elevated ArcA phosphorylation state under anaerobic conditions may explain the higher conjugation rate observed in IncF plasmids.

Among the *qnrS1*-IncF plasmids, the multireplicon IncFIA/FIB/FIC(FII) plasmid (pQNR278-15) achieved substantially higher transfer rates than the IncFIB/FIC(FII) plasmid (pQNR271-15) under anaerobic conditions, whereas no such differences was observed aerobically, suggesting specific plasmid or chromosome-encoded features in the anaerobic response. pQNR278-15 carries multiple *traN* copies and encodes anti-RM systems. TraN mediates mating-pair stabilisation through interactions with recipient outer membrane proteins. Multiple copies may enhance stabilisation under anaerobic conditions, where membrane composition or dynamics differ from aerobic conditions (Frankel et al., 2023; Low et al., 2022; Unden & Bongaerts, 1997).

The anti-RM systems on pQNR278-15 likely further increase transfer success (Dimitriu et al., 2024). The plasmid pQNR274-77 did not transfer to K12-*yfp* successfully under either condition. Although the three *qnrS1*-IncF plasmids shared a high degree of synteny, the differences between them may have been sufficient for the RM or CRISPR-Cas defence systems present in the K12-*yfp* to selectively target pQNR274-77, but not the remaining *qnrS1*-IncF plasmids (Garneau et al., 2010; Shaw et al., 2023).

Plasmids from other incompatibility groups have also shown contrasting oxygen responses. IncP-1β plasmids have been shown to have reduced transfer under anaerobic conditions (Król et al., 2011). Similarly, Ochi et al. (2021) reported a ∼10-fold decrease in IncP-1 conjugation frequency under anaerobic conditions in filter matings using bacteria from environmental samples. Oxygen-dependent patterns have also been described for other HGT mechanisms. It has been recently shown that anaerobic conditions increase cell-to-cell plasmid transformation frequency in *E. coli* biofilms (Hirayama et al., 2025). However, these studies did not account for changes in mating opportunities during the assays.

Our study offers several methodological advances over previous work. First, the use of the Approximate Extended Simonsen Model allowed us to decouple the conjugation rate from growth dynamics; an important consideration for our aerobic versus anaerobic comparisons, which likely affect (changes in) donor and recipient densities, hence mating opportunities (Huisman et al., 2022b; Shafieifini et al., 2022). Second, genomic characterisation of the recipients and donors enabled correlation with phenotypic conjugation outcomes, suggesting possible explanations for the observed patterns. Third, the fully factorial conjugation design using commensal *E. coli* donors and recipients carrying clinically relevant antimicrobial-resistant determinants allowed us to disentangle the influence of donor and recipient strain variation on plasmid transfer rate.

Although genomic characterisation identified candidate mechanisms for the observed variation in plasmid transfer rates, we did not experimentally link these features to oxygen-response outcomes. Transcriptomic or proteomic comparisons of donors and recipients under aerobic versus anaerobic conditions could reveal the regulatory mechanisms underlying plasmid and recipient-specific oxygen responses. Mutagenesis of candidate transfer genes could then establish causal relationships between plasmid genetic content and oxygen-dependent transfer phenotypes. Furthermore, our binary comparison may overlook threshold-dependent shifts in metabolic activity or regulatory ‘switches’ that may specifically occur under microaerobic conditions.

Our findings are also limited by the narrow range of plasmids, resistance genes, and strains examined. We focused on three natural variants of two plasmid types (IncI1-αand IncF) and two resistance genes (*bla*_CTX-M-1_ and *qnrS1*). While epidemiologically important, they represent only a small fraction of the plasmid diversity circulating in *E. coli* populations. The extent to which our findings may be generalised to other plasmid incompatibility groups remains to be determined. Additionally, *qnrS1*-IncF donors were tested only with K12-*yfp*, as attempts at commensal-to-commensal transfer were unsuccessful, leaving this ecologically relevant transfer context uncharacterised. Finally, our *in vitro* conditions do not fully match the complex large intestine environment, where short-chain fatty acids, mucus, host immune factors, and competing microbiota in a structured environment presenting heterogeneous conditions and local interactions may all modulate conjugation.

In conclusion, our findings suggest that conjugation-mediated AMR transfer may be underestimated when assessed under aerobic conditions. ESBL-IncI1-α and qnrS1-IncF plasmids transferred more efficiently under anaerobic conditions, indicating enhanced dissemination potential in the gut and other anaerobic environments. AMR studies should therefore use experimental conditions that closely reflect the relevant physiological and ecological niche.

## METHODS

### Bacterial strains

We conducted conjugation experiments using nine commensal *E. coli* strains previously isolated from the caeca of healthy chickens, as part of the Dutch national monitoring of antimicrobial resistance in animals (MARAN, 2019, 2023). In addition, the laboratory strain K12-*yfp* was included. Isolates were stored in 80% glycerol at −80 °C (Table S5).

**Donor strains**: Three *E. coli* isolates carried IncI1-α plasmids encoding the *bla*_CTX-M-1_ gene (pESBL33-68, pESBL34-08, and pESBL34-22), conferring cefotaxime resistance. Three additional *E. coli* isolates harboured multi-replicon IncF plasmids carrying a *qnrS1* gene (pQNR271-15, pQNR274-77, and pQNR278-15), conferring ciprofloxacin resistance.

**Recipient strains:** Three *E. coli* isolates carried chromosomal mutations conferring ciprofloxacin resistance. The fourth recipient was the *E. coli* K12 laboratory strain, which contained a yellow fluorescent protein (*yfp*) gene insertion and chloramphenicol resistance.

### Conjugation experimental design

Conjugation experiments were performed following the method described by (Huisman et al., 2022) with minor modifications as described below. All bacterial strains used in this study are listed in Table 1.

A full factorial design was implemented with three ESBL-carrying donors (33-68, 34-08, and 34-22) each paired with three ciprofloxacin resistant recipients (208-10, 259-14, and 270-23), resulting in nine donor-recipient combinations (Table S6). In addition, both ESBL and *qnrS1* donors were mated with the laboratory strain K12-*yfp,* which served as a universal reference recipient. This design enabled statistical control of donor plasmid and recipient genetic background effects in downstream analyses.

Conjugation experiments were performed under both aerobic and anaerobic environments, with three biologically independent replicates carried out on separate days using freshly prepared cultures.

**ESBL donor pairs**: Donors were selected on lysogeny agar (LA) supplemented with cefotaxime (1 mg/L). Recipients R1 (208-1), R2 (259-14) and R3 (270-23) were selected on LA supplemented with ciprofloxacin (2 mg/L), and K12-*yfp* on LA with chloramphenicol (50 mg/L).

***qnrS1* donor pairs**: Donors were selected on LA supplemented with ciprofloxacin (0.125 mg/L) and recipient K12-*yfp* on LA with chloramphenicol (50 mg/L).

Transconjugants from ESBL and *qnrS1* donor pairs were selected on LA using the respective double-antibiotic combinations.

### Conjugation assay

A single colony forming unit (CFU) from each strain was inoculated into Lysogeny broth (LB) and cultured overnight at 41 °C with shaking (200 rpm) under either aerobic or anaerobic conditions using a CLARIOstar microplate reader (BMG LABTECH, Ortenberg, Germany), equipped with an atmospheric control unit. For anaerobic cultures, LB was flushed with nitrogen before inoculation.

Overnight cultures were diluted 1:100 in fresh LB and incubated at 41 °C with shaking (200 rpm) to mid-exponential phase (OD_600_ = 0.4; measured via a spectrophotometer (Biochrom WPA S1200+) after approximately 1-2 h). Monocultures were then diluted 1:100 again in LB to maintain exponential growth. Diluted monocultures were serially diluted and plated on LA containing the appropriate single selective antibiotic (cefotaxime, ciprofloxacin or chloramphenicol) to determine initial cell density (CFU/ml). Undiluted monocultures were also plated on double-selective LA plates as negative controls. All plates were incubated at 41 °C for 24 h either aerobically or anaerobically.

Diluted donor and recipient cultures were mixed at a 1:1 ratio and vortexed. Control monocultures were prepared by mixing each diluted culture 1:1 with LB to measure donor and recipient individual growth rates for further conjugation rate calculations. Aliquots (200 µL) were pipetted in triplicate into 96-well plates and incubated statically at 41 °C in the CLARIOstar plate reader, with optical density measurements recorded every 20 min. Incubation durations were 3 h for anaerobic cultures and 5 h for aerobic cultures, both corresponding to entry into the stationary phase as determined by preliminary growth curve analysis.

Following incubation, conjugation mixtures were serially diluted in LB and plated on LA containing single selective antibiotics to enumerate donors and recipients, and double-selective antibiotics to enumerate transconjugants. Cell densities (CFU/ml) were calculated from colony counts and dilution factors. Transconjugant counts were subtracted from single-selective plate counts to obtain accurate donor and recipient densities. Monoculture controls were plated on double-selective LA to confirm the absence of spontaneous resistance. All plates were incubated at 41 °C for 24 h either aerobically or anaerobically.

### Putative transconjugant confirmation

Six putative transconjugant colonies were selected from each biological replicate for each donor-recipient pair to verify successful plasmid transfer. Selected colonies were re-streaked onto double-selective LA plates containing appropriate antibiotics.

DNA extraction was performed using the boiling method. Two to three colonies were suspended in 100 µL of sterile Milli-Q water and heated at 100 °C for 10 min, followed by cooling on ice for 5 min. Samples were then centrifuged at 17,530 x *g* for 5 min, and the supernatant was transferred to a fresh tube and stored at −20 °C until PCR analysis.

Transconjugant confirmation strategies varied depending on the donor-recipient pair (primer details in Table S7). CTX-M primers were used to confirm the presence of the IncI1-α-*bla*_CTX-M-1_ plasmid (pESBL33-68, pESBL34-08 or pESBL34-22) in transconjugants P1-P12, and *qnrS1* primers to confirm the presence of the IncF-*qnrS1* plasmid (pQNR271-15, pQNR274-77 or pQNR278-15) in transconjugants P13 to P15. Additionally, transconjugants P10-P15 were tested with *yfp* primers to confirm that the K12-*yfp* recipient strain had successfully acquired the donor plasmid.

All PCR reactions were performed in a final volume of 25 µL containing 12.5 µL Biomix^TM^ Red (Meridian Bioscience), 1 µL template DNA (2 µL for CTX-M), 10 pmol each of forward and reverse primers (20 pmol each for CTX-M) and nuclease-free water to volume. Donor and recipient strains were included as positive and negative controls for all PCR assays. Amplified products were analysed by electrophoresis on 1% agarose gels (for *yfp*) or 2% agarose gels (for CTX-M and *qnrS1*).

Transconjugants P1 to P9 were additionally screened on triple-selective LA containing cefotaxime (1 mg/L), ciprofloxacin (2 mg/L) and tetracycline (16 mg/L). This combination selected for the recipient’s background resistance (tetracycline and ciprofloxacin) while selecting for the donor plasmid, providing phenotypic confirmation that the recipient had acquired the IncI1-α-bla_CTX-M-1_ plasmid.

### Growth rates

The growth rates of donors, recipients and transconjugants were used to estimate the conjugation rates with the Approximate Extended Simonsen Model (ASM) (Huisman et al., 2022). After transconjugants were confirmed, one transconjugant from each of the three biological replicate conjugation experiments was selected per donor-recipient pair. A single CFU of each transconjugant was inoculated in LB and grown overnight at 41°C in a 12-well plate in the CLARIOstar plate reader under either aerobic or anaerobic environments.

Overnight cultures were diluted 1:100 in fresh LB and incubated at 41°C with shaking at 200 rpm. When cultures reached mid-exponential phase (OD600=0.4, measured by spectrophotometry), they were diluted 1:100 in LB. Triplicate aliquots from each culture were transferred into a 96-well plate and incubated statically at 41°C in the CLARIOstar plate reader. Bacterial growth was monitored by measuring optical density every 20 min.

Maximum specific growth rates (h^-1^) were calculated using the growthrates v0.8.4 package in R v4.2.3 (Hall et al., 2014; R Core Team, 2025). For transconjugants, growth rates were estimated per strain by pooling data from all three technical replicates and three independent biological replicates. For recipient and donor strains, growth rates were calculated per strain by pooling data from the three technical and biological replicates collected during the conjugation experiments.

Donor and recipient growth rates were estimated from CFU counts obtained during the conjugation experiments using the equation (Kirchman, 2002):

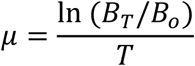

where *µ* is the specific donor or recipient growth rate, *B_T_* is the final donor or recipient population size, *B_0_* is the initial donor or recipient population size, and *T* is the conjugation duration (3 h for anaerobic and 5 h for aerobic experiments).

The transconjugant’s growth rate was corrected by multiplying the ratio of the transconjugant and recipient growth rates derived from OD₆₀₀ measurements by the recipient growth rate estimated from CFU counts, as described above.

### Conjugation rates estimation

Plasmid conjugation rates from donor–recipient mixed cultures were estimated using the Approximate Extended Simonsen Model (ASM) end-point formula implemented in the *conjugator* v.1.0.0 package (Huisman et al., 2022a). The ASM accounts for differences in donor and recipient growth rates during the mating period, providing a more accurate estimate of conjugation rates than traditional end-point methods. Conjugation rates were calculated for each independent biological replicate and log₁₀-transformed before statistical analysis.

### Statistical analysis

#### Full factorial experiment

For the 3×3×2 full factorial design, a linear mixed-effects model was fitted using the *lme4* v.1.1-38 package (Bates et al., 2015). Model assumptions were evaluated using diagnostic plots. Type III ANOVA with Satterthwaite’s approximation for denominator degrees of freedom was performed using the *lmerTest* v.3.1-3 package (Kuznetsova et al., 2017). Estimated marginal means and Tukey-adjusted pairwise contrasts were computed using the *emmeans* v.1.11.2-8 package. Statistical significance was defined as <0.05 for all tests. Data visualisation was performed using *ggplot2* v.3.5.2 (Wickham, 2016), and heatmaps were generated using the *ComplexHeatmap* v.2.25.2 package(Gu, 2022). All statistical analyses were performed in *R* v.4.5.1 (R Core Team, 2025a).

### DNA extraction and sequencing

Genomic DNA was extracted using the Puregene cell kit (Qiagen) following the manufacturer’s protocol for Gram-negative bacteria, with modifications (Qiagen, 2022). Briefly, bacterial colonies from overnight culture on LA plates were collected using a 10 µL loop and resuspended in 600 µL Cell Lysis Solution. Samples were mixed by pipetting and incubated at 85 °C for 5 min, then cooled on ice for 1 min.

RNA was removed by adding 3 µL of RNase A solution, mixing by inversion 25 times and incubating at 37 °C for 60 min. Samples were cooled on ice for 1 min, and proteins were separated by adding 200 µL Protein Precipitation Solution, vortexing for 20 s, and centrifuging at 13,000 x *g* for 3 min. The supernatant was transferred to a clean 1.5 ml tube containing 600 µL of isopropanol and mixed by inversion 50 times. DNA was pelleted by centrifugation at 13,000 x *g* for 1 min, washed with 600 µL of 70% ethanol, mixed by inversion 15 times and centrifuged at 13,000 x *g* for 1 min.

After discarding the supernatant, the pellet was air-dried for 15 min and resuspended in 100 µL of Elution Buffer (EB). DNA was dissolved by incubation at 56°C with gentle agitation for at least 1 h or until fully solubilised. DNA concentration and purity were assessed using the CLARIOstar, and samples were stored at −20 °C until sequencing.

Short-read sequencing libraries were prepared using the Illumina DNA prep kit, and paired-end sequencing was performed on either the NextSeq 2000 or NovaSeq 6000 Sequencing System (Illumina). Long-read sequencing libraries were prepared using the Native Barcoding Kit 96 V14 and sequenced on a PromethION instrument with a R10.4.1 flow cell (Oxford Nanopore Technologies) for 72 h. Basecalling was performed with Dorado v4.2.0 or v5.0.0 using the high-accuracy method.

All recipient strains were sequenced using short-read sequencing, whereas all the donor strains were sequenced using both short and long read methods to fully resolve the plasmids of interest.

### De novo assembly and annotation

Raw short-reads were trimmed using *BBTools suite* v39.26, and raw long-reads were trimmed using *Filtlong* v0.2.1 (Bushnell, 2014; R. Wick, 2017/2026). Read quality was assessed with *FastQC* v0.12.1 (FastQC, 2015). Genome assembly was performed using *Unicycler* v0.5.1 for both short-read assemblies and hybrid assemblies combining long and short reads (R. R. Wick et al., 2017). Assembly quality was assessed using *QUAST* v5.2.08 (Gurevich et al., 2013). Unicycler output was used to evaluate plasmid circularisation.

Genome annotation was carried out with *Bakkta* v1.8.1, with the full database v5.1 (Schwengers et al., 2021). Antimicrobial resistance genes, including chromosomal resistance-associated mutations, were identified using *ResFinder* v4.7.2 (Bortolaia et al., 2020), and plasmid replicons with *PlasmidFinder* v2.0.1 (Carattoli et al., 2014). Defence systems were detected with *DefenseFinde*r v2.0.1 (Couvin et al., 2018; Néron et al., 2023; Tesson et al., 2022). Multi-locus sequence types (MLST) were assigned using the Achtman scheme implemented in *mlst* v2.23.0 (Seemann, 2014/2025) and *E coli* phylogroups were determined using *ClermonTyping* v24.02 (Beghain et al., 2018).

### Core genome phylogenetic analysis

Pangenome analysis was performed using *Panaroo* v1.5.2, with GFF3 annotation files generated by Bakta as input (Tonkin-Hill et al., 2020). Maximum likelihood phylogenetic reconstruction was conducted using *IQ-TREE* v3.0.1 on the core genome alignment (Wong et al., 2025). The optimal nucleotide substitution model was selected using ModelFinder (Kalyaanamoorthy et al., 2017), and branch support was assessed using 1,000 ultrafast bootstrap replicates (Hoang et al., 2018). The resulting maximum likelihood tree was mid-point rooted and visualised in R v4.2.3 using *ggtree* v3.14.0 (R Core Team, 2025b; Yu et al., 2017).

### Plasmid analysis

The six plasmids of interest were confirmed to be circular based on the Unicycler output and were selected for further analysis. Plasmids pQNR271-15, pQNR274-77, and pQNR278-15 were analysed together, and plasmids pESBL33-68, pESBL34-08, and pESBL34-22 were analysed as a separate group. All plasmid sequences were first reoriented to begin at a conserved reference region identified through k-mer analysis (21-bp k-mers; minimum conserved threshold 0.8). Reoriented plasmid sequences were aligned using *progressiveMauve* v1.6.1 using default parameters to assess genome synteny (Darling et al., 2010). The resulting multiple-sequence alignment (XMFA) enabled subsequent identification of conserved genomic regions, structural variations (including inversions and translocations) and regions of differential gene content among the plasmids.

Integrated visualisation of gene annotations and sequence alignments was performed using the *genoPlotR* v0.8.11 package in R v4.2.3 (Guy et al., 2010; R Core Team, 2025b). Annotations generated by *Bakta* and *DefenseFinder* v2.0.1 were used for plasmid feature annotation (Schwengers et al., 2021; Tesson et al., 2022). The progressiveMauve backbone file (XMFA) was imported with a minimum gene size threshold of 400bp to reduce visualisation complexity.

## Acknowledgements

The authors thank Alieda van Essen and Michiel de Boer for their technical support and Frank Harders for his assistance with DNA sequencing. We also thank Benno ter Kuile and Nadine Handel (University of Amsterdam) for providing E.coli strain K12-*yfp*.

This research was funded by the Netherlands Ministry of Agriculture, Fisheries, Food Security and Nature under grant agreement KB-37-003-024 and the JPIAMR program STRESST, funded by ZonMw under grant agreement 10570132110004.

## Author contributions

IC-R conceptualised the study. IC-R, SF, AdV, MB and KV contributed to the design of the experiments. SF and IC-R performed the experiments. IC-R and SF analysed the data. IC-R and SF wrote the original draft of the manuscript. All authors reviewed and edited the final manuscript. MB acquired funding. MB, AdV and KV supervised the project.

## Data availability

The data supporting the findings of this study are available from the corresponding author upon reasonable request. The data will be deposited in an appropriate public repository upon publication.

