## Supplemental file 1 for "Anaerobic conditions increase plasmid transfer rates across *Escherichia coli* strains"

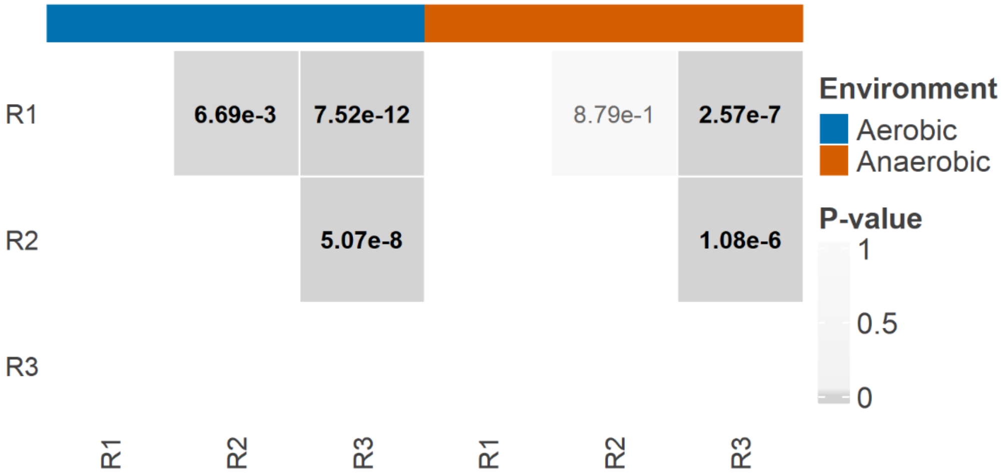


**Figure S1. Pairwise Tukey-adjusted comparison heatmap for QRDR recipients–environment combinations.** Cell shading represents *p*-values, with darker grey indicating lower values. Significant comparisons ($p \leq 0.05$) are shown in bold. The annotation bar indicates both environments.

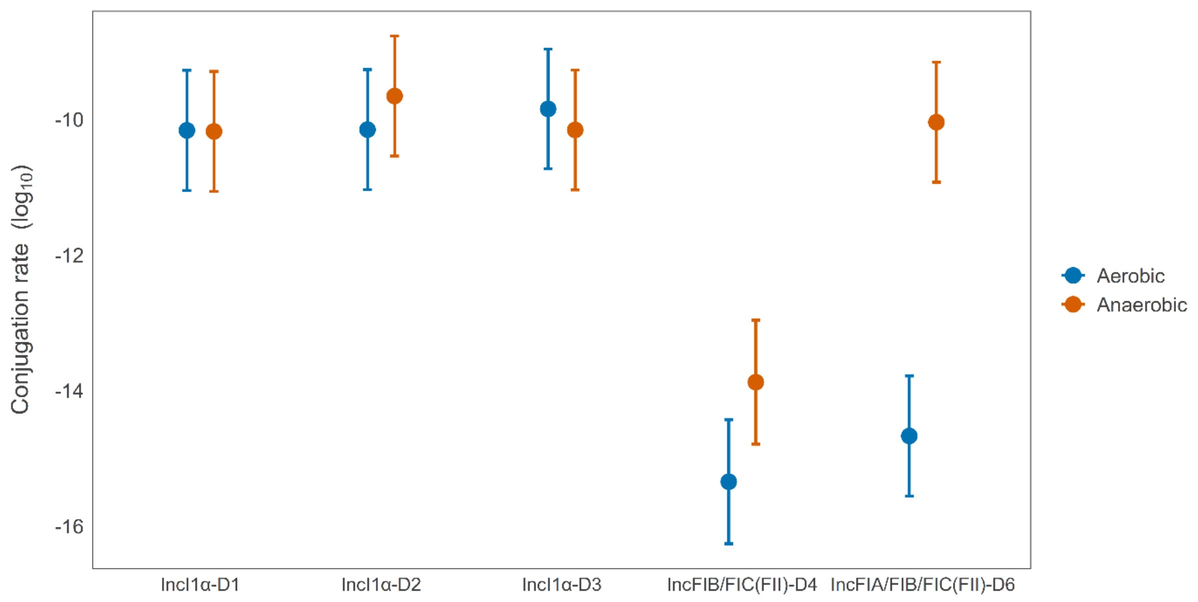


**Figure S2. Estimated marginal means of conjugation rates [mL (CFU h)−1] for plasmid type under aerobic and anaerobic conditions.** Dots denote means; error bars indicate 95% confidence intervals.

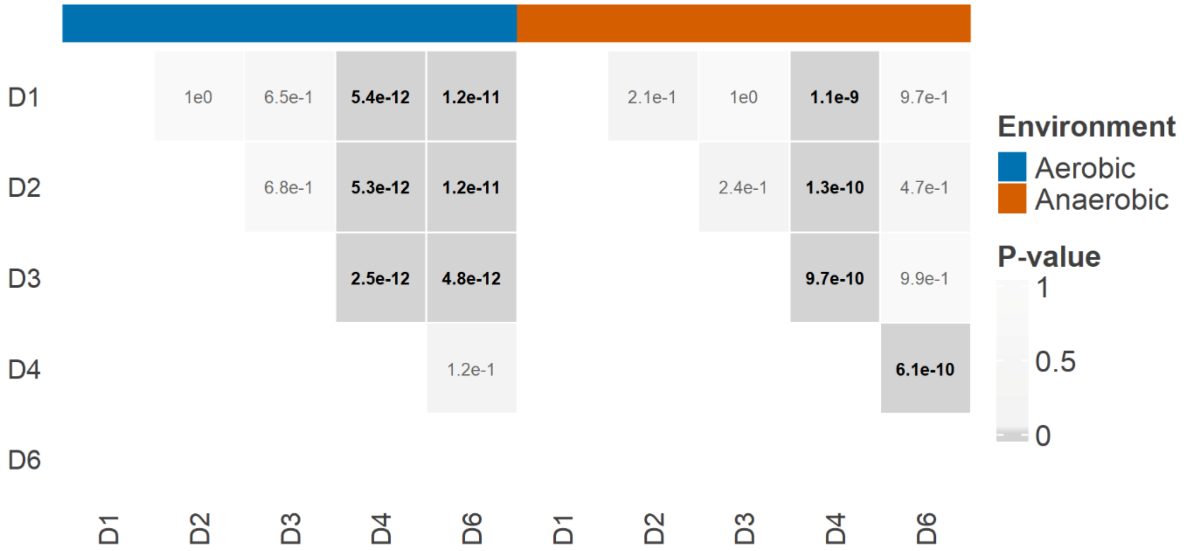


**Figure S3. Pairwise Tukey-adjusted comparison heatmap for ESBL and *qnr*-carrying donor–environment combinations**. Cell shading represents p-values, with darker grey indicating lower values. Significant comparisons ($p \leq 0.05$) are shown in bold. The annotation bar indicates both environments.

**Table S1. Effects of environment, donor, and recipient strain on ESBL-Inc1α conjugation rates.**

| Fixed effects | Estimate | SE[a] | t value | DF[b] | P value |
|---|---|---|---|---|---|
| (Intercept) | -14.09 | 0.20 | -68.79 | 34 | **1.7e-38** |
| Anaerobic environment | 0.66 | 0.28 | 2.34 | 34 | **0.025** |
| DonorD2 | -0.45 | 0.28 | -1.60 | 34 | 0.119 |
| Donor D3 | -0.07 | 0.28 | -0.24 | 34 | 0.814 |
| Recipient R2 | 0.42 | 0.28 | 1.48 | 34 | 0.144 |
| Recipient R3 | 1.59 | 0.28 | 5.65 | 34 | **2.5e-06** |
| Anaerobic environment x Donor D2 | 0.37 | 0.40 | 0.94 | 34 | 0.354 |
| Anaerobic environment x Donor D3 | -0.18 | 0.40 | -0.46 | 34 | 0.647 |
| Anaerobic environment x Recipient R2 | -0.28 | 0.40 | -0.71 | 34 | 0.483 |
| Anaerobic environment x Recipient R3 | -0.43 | 0.40 | -1.09 | 34 | 0.284 |
| Donor D2 x Recipient R2 | 0.60 | 0.40 | 1.50 | 34 | 0.142 |
| Donor D3 x Recipient R2 | -0.26 | 0.40 | -0.67 | 34 | 0.509 |
| Donor D2 x Recipient R3 | 0.17 | 0.40 | 0.42 | 34 | 0.675 |
| Donor D3 x Recipient R3 | 0.23 | 0.40 | 0.58 | 34 | 0.563 |
| Anaerobic environment x Donor D2 x Recipient R2 | -0.96 | 0.56 | -1.70 | 34 | 0.099 |
| Anaerobic environment x Donor D3 x Recipient R2 | 0.44 | 0.56 | 0.79 | 34 | 0.437 |
| Anaerobic environment x Donor D2 x Recipient R3 | -0.69 | 0.56 | -1.22 | 34 | 0.229 |
| Anaerobic environment x Donor D3 x Recipient R3 | 0.12 | 0.56 | 0.21 | 34 | 0.832 |

| Group | Random effects | Variance | SD[c] |
|---|---|---|---|
| Independent biological replicates | Intercept | 0.006 | 0.080 |
| Residual | | 0.119 | 0.346 |

Linear mixed-effects model fitted to $\log_{10}$-transformed conjugation rates from the full factorial experiment (3 ESBL donors × 3 QRDR recipients × 2 environments). Fixed effects included environment (aerobic vs. anaerobic), donor strain, recipient strain, and all two-way and three-way interactions. A random intercept for experimental day was included to account for temporal variation across independent replicates. Significant *p*-values (*p* <0.05) are shown in bold.

[a]SE, standard error.

[b]Degrees of freedom calculated using Satterthwaite's approximation.

[c]SD, standard deviation.

**Table S2. Effects of oxygen availability and plasmid type on conjugation efficiency.**

| Fixed effects | Estimate | SE[a] | t value | DF[b] | P value |
|---|---|---|---|---|---|
| Intercept | -10.16 | 0.45 | -22.53 | 18 | 1.2e-14 |
| Anaerobic environment | -0.02 | 0.23 | -0.07 | 18 | 0.948 |
| Plasmid type (Inc1α-D2) | 0.01 | 0.64 | 0.02 | 18 | 0.986 |
| Plasmid type (Inc1α-D3) | 0.32 | 0.64 | 0.50 | 18 | 0.626 |
| Plasmid type (FIA_IncFIB-D4) | -5.18 | 0.65 | -8.00 | 18 | **2.5e-07** |
| Plasmid type (IncFIB_IncFIC-D6) | -4.51 | 0.64 | -7.07 | 18 | **1.4e-06** |
| Anaerobic environment x Plasmid type (Inc1α-D2) | 0.51 | 0.33 | 1.56 | 18 | 0.136 |
| Anaerobic environment x Plasmid type (Inc1α-D3) | -0.30 | 0.33 | -0.90 | 18 | 0.379 |
| Anaerobic environment x Plasmid type (FIA_IncFIB-D4) | 1.48 | 0.37 | 4.05 | 18 | **7.4e-04** |
| Anaerobic environment x Plasmid type (IncFIB_IncFIC-D6) | 4.64 | 0.33 | 14.18 | 18 | **3.3e-11** |

| Group | Random effects | Variance | SD[c] |
|---|---|---|---|
| Donor | (Intercept) | 0.177 | 0.420 |
| Residual | | 0.080 | 0.283 |

Linear mixed-effects model fitted to $log_{10}$-transformed conjugation rates with environment (aerobic vs anaerobic) and plasmid type as fixed effects, their two-way interactions, and donor strain as random intercepts to control for donor biological variation. Significant *p*-values ($p < 0.05$) are shown in bold.

[a]SE, standard error.

[b]Degrees of freedom calculated using Satterthwaite's approximation.

[c]SD, standard deviation.

**Table S3. Effects of oxygen availability and donor strain on conjugation efficiency.**

| Term | SS[a] | MS[b] | $DF_{Num.Den}$[c] | F[d] | P value | Explained variance (%)[e] |
|---|---|---|---|---|---|---|
| Environment | 10.71 | 10.71 | 1, 16.3 | 134.87 | **2.6e-09** | 9.21 |
| Donor | 79.26 | 19.82 | 4, 16.6 | 249.63 | **1.3e-14** | 68.16 |
| Environment x Donor | 24.07 | 6.02 | 4, 16.3 | 75.80 | **2.5e-10** | 20.69 |
| Residuals | | | | | | 1.91 |

| Group | Random Effect | Variance | SD[f] |
|---|---|---|---|
| Day | (Intercept) | 0.001 | 0.031 |
| Residual | | 0.079 | 0.282 |

| Fixed effects | Estimate | SE[g] | t value | DF | P value |
|---|---|---|---|---|---|
| Intercept | -10.16 | 0.16 | -62.09 | 17.98 | **2.0e-22** |
| Anaerobic environment | -0.02 | 0.23 | -0.07 | 16.32 | 0.948 |
| Donor D2 | 0.01 | 0.23 | 0.05 | 16.32 | 0.961 |
| Donor D3 | 0.32 | 0.23 | 1.37 | 16.32 | 0.188 |
| Donor D4 | -5.18 | 0.26 | -20.13 | 16.84 | **3.2e-13** |
| Donor D6 | -4.51 | 0.23 | -19.60 | 16.32 | **9.1e-13** |
| Anaerobic environment x Donor D2 | 0.51 | 0.32 | 1.57 | 16.32 | 0.136 |
| Anaerobic environment x Donor D3 | -0.29 | 0.32 | -0.91 | 16.32 | 0.378 |
| Anaerobic environment x Donor D4 | 1.48 | 0.36 | 4.08 | 16.32 | **8.4e-04** |
| Anaerobic environment x Donor D6 | 4.64 | 0.32 | 14.27 | 16.32 | **1.2e-10** |

Type III ANOVA and linear mixed-effects model fitted to $\log_{10}$-transformed conjugation rates with environment (aerobic vs anaerobic) and donor strain as fixed effects, their two-way interactions, and experimental day as random intercepts to control for day-to-day biological variation. Degrees of freedom were calculated using Satterthwaite's approximation. Significant *p*-values ($p < 0.05$) are shown in bold.

[a]SS, sum of squares.

[b]MS, mean squares.

[c]$DF_{Num, Den}$, numerator and denominator degrees of freedom calculated using Satterthwaite's approximation.

[d]F, F-statistic.

[e]Explained variance (%), proportion of the total sum of squares for each fixed term and residual.

[f]SD, standard deviation.

[g]SE, standard error.

**Table S4. Fold-change in conjugation rates between aerobic and anaerobic conditions for individual donors.**

| **Contrast**[a] | **Donor** | **Ratio**[b] | **SE**[c] | **DF**[d] | **Null** | **t ratio**[e] | **P value** |
|---|---|---|---|---|---|---|---|
| Aerobic / Anaerobic | D1 | 1.04 | 0.55 | 16 | 1 | 0.067 | 0.948 |
| Aerobic / Anaerobic | D2 | 0.32 | 0.17 | 16 | 1 | -2.152 | **0.047** |
| Aerobic / Anaerobic | D3 | 2.04 | 1.08 | 16 | 1 | 1.349 | 0.196 |
| Aerobic / Anaerobic | D4 | 0.03 | 0.02 | 16 | 1 | -5.212 | **8.52e-05** |
| Aerobic / Anaerobic | D6 | 2.36e-05 | 1.25e-05 | 16 | 1 | -20.115 | **8.56e-13** |

[a]Pairwise comparison for ESBL (D1-D3) and *qnrS1* (D4 and D6) donors. Significant *p*-values ($p < 0.05$) are shown in bold.

[b]Ratio, *fold-change* between aerobic and anaerobic conditions.

[c]SE, standard error.

[d]Degrees of freedom calculated using the Kenward-Roger method.

[e]t ratio, test statistic used to assess whether the Aerobic/Anaerobic ratio differs from the null value of 1.

**Table S5. Genomic characteristics of donor and recipient isolates.**

| Sample ID | Type[a] | MLST[b] | Phylogroup | Resistance genes in isolate | Defence genes in isolate | Plasmid replicons in isolate |
|---|---|---|---|---|---|---|
| 33-68 | D1 | 101 | B1 | *aadA5*, *bla*CTX-M-1, *sul2*, *dfrA17* | cas2_I-E_2, cas1_I-E_1, cas6e_I_II_III_IV_V_VI_1, cas5_I-E_3, cas7_I-E_2, cse2gr11_I-E_1, cas8e_I-E_1, cas3_I_5, MazEF__MazF, MazEF__MazE, Druantia_I__DruA, Druantia_I__DruB, Druantia_I__DruC, Druantia_I__DruD, Druantia__DruE_1, MazEF__MazE, MazEF__MazF, CBASS__A_Cyclase_AGS_C, CBASS__2TM_Gros, Gao_Qat__QatD, Gao_Qat__QatC, Gao_Qat__QatB, Gao_Qat__QatA, PD-Lambda-1__PD-Lambda-1, klca, MazEF__MazE, MazEF__MazF, RM_Type_IV__Type_IV_01, RM_Type_IV__FAM_1, RM_Type_IV__Type_IV_22, RM__Type_I_S_52, RM__Type_I_MTases_FAM_1, RM__Type_I_REases_FAM_2.einsi_trimmed, DRT8__DRT8a, DRT8__DRT8b, DarTG__DarT, DarTG__DarG, TIR-IV__TIR-IV_A, TIR-IV__TIR-IV_B, racc, Rst_3HP__Hp1, Rst_3HP__Hp2, Rst_3HP__Hp3, Lamassu-Fam__LmuB_SMC_Lipase, Lamassu-Fam__LmuC_acc_Lipase, Lamassu-Fam__LmuA_effector_Lipase, ral, abc2, ardb, psia, psib, arda, TIR-III__TIR-III_A, TIR-III__TIR-III_B, Pif__PifA, Pif__PifC, ardb, psia, psib, Mok_Hok_Sok__Mok | Col(MG828), Col156, IncFIA/IncFIC(FII), IncI1-I(Alpha) |
| 34-08 | D2 | 155 | B1 | *bla*CTX-M-1, *sul2* | RM_Type_IV__Type_IV_05, SanaTA__SanaT, SanaTA__SanaA, PrrC__EcoprrI, RM__Type_I_S_01, RM__Type_I_REases_FAM_0.einsi_trimmed, MazEF__MazF, MazEF__MazE, SDIC3__SDIC3F, SDIC3__SDIC3E, SDIC3__SDIC3C, MazEF__MazE, MazEF__MazF, cas3_I_5, cas8e_I-E_1, cse2gr11_I-E_1, cas7_I-E_2, cas5_I-E_3, cas6e_I_II_III_IV_V_VI_1, cas1_I-E_1, cas2_I-E_2, PD-Lambda-4__PD-Lambda-4_B, PD-Lambda-4__PD-Lambda-4_A, abc2, RM_Type_IIG__Type_IIG_FAM_1.einsi_trimmed, klca, racc, gam, PD-T7-1__PD-T7-1, RM_Type_II__Type_II_REase27, RM_Type_II__Type_II_MTases_FAM_2, PD-T4-3__PD-T4-3, Mok_Hok_Sok__Mok, psib, psia, ardb, ardb, psia, psib, arda | Col(MG828), ColpVC, IncFIA/IncFIB(AP001918)/IncFIC(FII), IncI1-I(Alpha) |
| 34-22 | D3 | 101 | B1 | *bla*CTX-M-1, *sul2*, *dfrA17* | MazEF__MazE, MazEF__MazF, RM_Type_IV__Type_IV_01, RM_Type_IV__FAM_1, RM_Type_IV__Type_IV_22, RM__Type_I_S_52, RM__Type_I_MTases_FAM_1, RM__Type_I_REases_FAM_2.einsi_trimmed, DRT8__DRT8a, DRT8__DRT8b, Lamassu-Fam__LmuA_effector_Lipase, Lamassu-Fam__LmuC_acc_Lipase, | Col156, IncFIA/IncFIC(FII), IncI1-I(Alpha) |

| | | | | | | |
|---|---|---|---|---|---|---|
| | | | | | Lamassu-Fam__LmuB_SMC_Lipase, Rst_3HP__Hp3, Rst_3HP__Hp2, Rst_3HP__Hp1, racc, TIR-IV__TIR-IV_B, TIR-IV__TIR-IV_A, DarTG__DarG, DarTG__DarT, klca, PD-Lambda-1__PD-Lambda-1, Gao_Qat__QatA, Gao_Qat__QatB, Gao_Qat__QatC, Gao_Qat__QatD, CBASS__2TM_Gros, CBASS__A_Cyclase_AGS_C, MazEF__MazF, MazEF__MazE, Druantia__DruE_1, Druantia_I__DruD, Druantia_I__DruC, Druantia_I__DruB, Druantia_I__DruA, MazEF__MazE, MazEF__MazF, cas3_I_5, cas8e_I-E_1, cse2gr11_I-E_1, cas7_I-E_2, cas5_I-E_3, cas6e_I_II_III_IV_V_VI_1, cas1_I-E_1, cas2_I-E_2, ardb, psia, psib, arda, TIR-III__TIR-III_A, TIR-III__TIR-III_B, Pif__PifA, Pif__PifC, ardb, psia, psib, Mok_Hok_Sok__Mok | |
| 271-15 | D4 | 162 | B1 | *bla*TEM-135, *qnrS1*, *tet(A)* | klca, MazEF__MazE, MazEF__MazF, cas3_I_5, cas8e_I-E_1, cse2gr11_I-E_1, cas7_I-E_2, cas5_I-E_3, cas6e_I_II_III_IV_V_VI_1, klca, Kiwa__KwaA, Kiwa__KwaB, klca, BstA__BstA, gam, DS-17__DS-17, klca, CBASS__2TM_new, CBASS__Cyclase_SMODS, DS-13__DS-13A, DS-13__DS-13B, DS-13__DS-13B, MazEF__MazF, MazEF__MazE, Lamassu-Fam__LmuB_SMC_Mrr, Lamassu-Fam__LmuC_acc_Mrr, Lamassu-Fam__LmuA_effector_Mrr, MazEF__MazF, MazEF__MazE, Mok_Hok_Sok__Mok, psib, psia, ardb, psia, psib, arda | ColpVC, IncFIB(AP001918)/IncFIC(FII), IncI1-I(Alpha) |
| 274-77 | D5 | 162 | B1 | *bla*TEM-135, *qnrS1*, *tet(A)* | klca, MazEF__MazE, MazEF__MazF, cas3_I_5, cas8e_I-E_1, cse2gr11_I-E_1, cas7_I-E_2, cas5_I-E_3, cas6e_I_II_III_IV_V_VI_1, cas1_I-E_1, cas2_I-E_2, klca, Kiwa__KwaA, Kiwa__KwaB, BstA__BstA, gam, DS-17__DS-17, klca, CBASS__2TM_new, CBASS__Cyclase_SMODS, DS-13__DS-13A, DS-13__DS-13B, DS-13__DS-13B, MazEF__MazF, MazEF__MazE, Dnd__DndB, Dnd__DndC, Dnd__DndD, Dnd__DndE, Dnd_ABCDEFGH__DptH, Dnd_ABCDEFGH__DptG, Dnd_ABCDEFGH__DptF, klca, Lamassu-Fam__LmuB_SMC_Mrr, Lamassu-Fam__LmuC_acc_Mrr, Lamassu-Fam__LmuA_effector_Mrr, MazEF__MazE, MazEF__MazF, Mok_Hok_Sok__Mok, psib, psia | IncFIB(AP001918)/IncFIC(FII) |
| 278-15 | D6 | 1485 | F | *aph(6)-Id*, *aph(3'')-Ib*, *bla*TEM-1B, *qnrS1*, *sul2*, *tet(A)* | PsyrTA__PsyrT, PsyrTA__PsyrA, MazEF__MazE, MazEF__MazF, cas3_I_5, cas8e_I-E_1, cse2gr11_I-E_1, cas7_I-E_2, cas5_I-E_3, cas6e_I_II_III_IV_V_VI_1, cas1_I-E_1, cas2_I-E_2, Lamassu-Fam__LmuB_SMC_Cap4_nuclease_II, Lamassu-Fam__LmuC_acc_Cap4_nuclease, Lamassu-Fam__LmuA_effector_Cap4_nuclease_II, Septu__PtuB, Septu__PtuA, | Col(MG828), Col156, IncFIA/IncFIB(AP001918)/IncFIC(FII), IncFII(pSE11), IncI2(Delta), IncP1 |

| | | | | | | |
|---|---|---|---|---|---|---|
| | | | | | RM__Type_I_MTases_FAM_0, RM__Type_I_S_04, RM__Type_I_REases_FAM_0.einsi_trimmed, racc, gam, RM__Type_I_REases_FAM_2.einsi_trimmed, RM__Type_I_MTases_FAM_2, RM__Type_I_S_52, RM_Type_IV__FAM_1, RM_Type_IV__FAM_2, Druantia__DruE_3, Druantia_III__DruH, klca, MazEF__MazF, MazEF__MazE, DS-17__DS-17, BREX__brxA_DUF1819, BREX__brxB_DUF1788, BREX__brxC, BREX__pglX1, BREX__pglZA, BREX__brxL, Dnd_ABCDEFGH__DptF, Dnd_ABCDEFGH__DptG, Dnd_ABCDEFGH__DptH, Dnd__DndE, Dnd__DndD, Dnd__DndC, Dnd__DndB, klca, Mok_Hok_Sok__Mok, psib, psia, ardb, DS-28__DS-28B, DS-28__DS-28A, Gao_Tmn__TmnA, ardb, psia, psib, Mok_Hok_Sok__Mok, aca5, Mok_Hok_Sok__Mok, ardk, klca, ardc | |
| 208-1 | R1 | 155 | B1 | *bla*TEM-1B, *catA1*, *tet(B),* gyrA p.S83L, gyrA p.D87N, parC p.S80I | Shango__SngA, Shango__SngB, Shango__SngC, DS-17__DS-17, BREX__brxA_DUF1819, BREX__brxB_DUF1788, BREX__brxC, BREX__pglX1, Dpd__DpdC, Dpd__DpdA, Dpd__DpdB, Dpd__DpdD, Dpd__DpdK, Dpd__DpdJ, Dpd__DpdI, Dpd__DpdH, Dpd__DpdG, Dpd__DpdF, Dpd__DpdE, Dpd__QueD, MazEF__MazF, MazEF__MazE, racc, MazEF__MazF, MazEF__MazE, klca, Dnd__DndB, Dnd__DndC, Dnd__DndD, Dnd__DndE, Dnd_ABCDEFGH__DptH, Dnd_ABCDEFGH__DptG, Dnd_ABCDEFGH__DptF, racc, mom, cas2_I-E_2, cas1_I-E_1, cas6e_I_II_III_IV_V_VI_1, cas5_I-E_3, cas7_I-E_2, cse2gr11_I-E_2, cas8e_I-E_1, cas3_I_5, PD-Lambda-4__PD-Lambda-4_B, PD-Lambda-4__PD-Lambda-4_A, MazEF__MazF, MazEF__MazE, RM_Type_IV__FAM_2, RM_Type_IV__FAM_1, Mokosh_TypeII__MkoC, RM__Type_I_REases_FAM_0.einsi_trimmed, RM__Type_I_S_51, PrrC__PrrC, RM__Type_I_MTases_FAM_0, psia, psib, Mok_Hok_Sok__Mok, MazEF__MazF, MazEF__MazE | IncFIB(AP001918), IncFIC(FII), p0111 |
| 259-14 | R2 | 162 | B1 | *bla*TEM-1B, *tet(A),* gyrA p.S83L, gyrA p.D87N, parC p.S80I | DS-17__DS-17, klca, Kiwa__KwaA, Kiwa__KwaB, Kiwa__KwaB_2, Kiwa__KwaA, RM__Type_I_REases_FAM_1.einsi_trimmed, RM__Type_I_S_52, RM__Type_I_MTases_FAM_3, MazEF__MazF, MazEF__MazE, klca, DS-29__DS-29, RM__Type_I_MTases_FAM_0, RM__Type_I_S_51, DarTG__DarT, DarTG__DarG, RM__Type_I_REases_FAM_0.einsi_trimmed, Mokosh_type_I__MkoB_A, Mokosh_type_I__MkoA_A, klca, MazEF__MazE, MazEF__MazF, cas3_I_5, cas8e_I-E_1, cse2gr11_I-E_1, cas7_I-E_2, cas5_I-E_3, cas6e_I_II_III_IV_V_VI_1, cas1_I-E_1, cas2_I-E_2, Kiwa__KwaB, Kiwa__KwaA, Gao_Hhe__HheA, TgvAB__TgvB, TgvAB__TgvA, RM__Type_I_REases_FAM_0.einsi_trimmed, RM__Type_I_S_51, | Col(pHAD28), IncFIB(AP001918), IncFIC(FII), IncY |

| | | | | | | |
|---|---|---|---|---|---|---|
| | | | | | RM__Type_I_MTases_FAM_0, Retron_XII__RT_12, gam, BstA__BstA, darb, ulx, PrrC__EcoprrI, RM__Type_I_S_01, RM__Type_I_REases_FAM_0.einsi_trimmed, hdf, dara, ddra, ddrb, dGTPase__Sp_dGTPase, MazEF__MazE, MazEF__MazF, klca, Mok_Hok_Sok__Mok, psib, psia | |
| 270-23 | R3 | 162 | B1 | *bla*TEM-1B, *catA1*, *tet(B),* gyrA p.S83L, gyrA p.D87N, parC p.S80I | klca, MazEF__MazE, MazEF__MazF, cas3_I_5, cas8e_I-E_1, cse2gr11_I-E_1, cas7_I-E_2, cas5_I-E_3, cas6e_I_II_III_IV_V_VI_1, cas1_I-E_1, cas2_I-E_2, DS-17__DS-17, klca, SDIC4__SDIC4B, SDIC4__SDIC4A, Dpd__QueD, Dpd__DpdD, Dpd__DpdK, Dpd__DpdJ, Dpd__DpdI, Dpd__DpdH, Dpd__DpdG, Dpd__DpdF, Dpd__DpdE, Dpd__DpdB, Dpd__DpdA, Dpd__DpdC, MazEF__MazF, MazEF__MazE, klca, Kiwa__KwaA, Kiwa__KwaB, gam, BstA__BstA, PD-T7-5__PD-T7-5, rad, MazEF__MazF, MazEF__MazE, ardb, psia, psib, arda, MazEF__MazE, MazEF__MazF, psia, psib, Mok_Hok_Sok__Mok | IncB/O/K/Z, IncFIB(AP001918), IncFIB(H89-PhagePlasmid), IncFIC(FII) |
| K12-yfp | R4 | 10 | A | *catA1* | MazEF__MazF, MazEF__MazE, Hachiman__HamB, Hachiman__HamA_2, RnlAB__RnlA, RnlAB__RnlB, klca, cas2_I-E_2, cas1_I-E_1, cas6e_I_II_III_IV_V_VI_1, cas5_I-E_3, cas7_I-E_2, cse2gr11_I-E_1, cas8e_I-E_1, cas3_I_5, MazEF__MazF, MazEF__MazE, RM_Type_IV__FAM_2, RM_Type_IV__FAM_1, RM__Type_I_S_52, RM__Type_I_MTases_FAM_2, RM__Type_I_REases_FAM_2.einsi_trimmed, Lit__Lit, RM_Type_IV__Type_IV_05, racc, klca | -[c] |

[a]Genomic features of ten *Escherichia coli* isolates. D1-D3, extended-spectrum beta-lactamase (ESBL) donors (*bla*$_{CTX-M-1}$); D4-D6, QNR donors (*qnrS1*), R1-R3, Quinolone Resistance-Determining Region (QRDR) recipients (chromosomal mutations in gyrA and parC), and a K12-yfp laboratory recipient.

[b]Multilocus sequence type (MLST)

[c]A dash indicates no plasmid replicons detected.

**Table S6. Donor–recipient mating pairs used in this study and their corresponding plasmids.**

| Name of transconjugant | Donor | Recipient | Plasmid |
|---|---|---|---|
| P1 | D1 (33-68) | R1 (208-10) | pESBL33-68 |
| P2 | D1 (33-68) | R2 (259-14) | pESBL33-68 |
| P3 | D1 (33-68) | R3 (270-23) | pESBL33-68 |
| P4 | D2 (34-08) | R1 (208-10) | pESBL34-08 |
| P5 | D2 (34-08) | R2 (259-14) | pESBL34-08 |
| P6 | D2 (34-08) | R3 (270-23) | pESBL34-08 |
| P7 | D3 (34-22) | R1 (208-10) | pESBL34-22 |
| P8 | D3 (34-22) | R2 (259-14) | pESBL34-22 |
| P9 | D3 (34-22) | R3 (270-23) | pESBL34-22 |
| P10 | D1 (33-68) | R4 (K12-*yfp*) | pESBL33-68 |
| P11 | D2 (34-08) | R4 (K12-*yfp*) | pESBL34-08 |
| P12 | D3 (34-22) | R4 (K12-*yfp*) | pESBL34-22 |
| P13 | D4 (271-15) | R4 (K12-*yfp*) | pQNR271-15 |
| P14 | D5 (274-77) | R4 (K12-*yfp*) | pQNR274-77 |
| P15 | D6 (278-15) | R4 (K12-*yfp*) | pQNR278-15 |

P1–P9 (highlighted in grey) represent the donor–recipient combinations included in the full factorial analysis.

**Table S7. PCR assay parameters**

| Primer | Sequence(5'-3') | Size of product (bp) | PCR conditions | Reference |
|---|---|---|---|---|
| *yfp*-F | TGAAGTCAGCCCCATACGAT | 1999 | 95 °C 10 min; (95 °C 15 s, 59 °C 1 min, 72 °C 30 s) × 30; 72 °C 7 min | (Elowitz et al., 2002) |
| *yfp*-R | GAGTCAGTGAGCGAGGAAGC | | | |
| *qnrS*-F | CGACGTGCTAACTTGCGTGATA | 538 | 96 °C 5 min; (94 °C 1 min, 57 °C 1 min, 72 °C 30 s) × 30; 72 °C 10 min | (Cavaco et al., 2008) |
| *qnrS*-R | TACCCAGTGCTTCGAGAATCAG | | | |
| CTX-M-F | ATGTGCAGYACCAGTAARGTKATGGC | 592 | 94 °C 5 min; (94 °C 30 s, 55 °C 30 s, 72 °C 1 min) × 30; 72 °C 5 min | (Mulvey et al., 2004) |
| CTX-M-R | TGGGTRAARTARGTSACCAGAAYCAGCGG | | | |

PCR primers, expected amplicon sizes, and amplification conditions used to confirm the presence *of yfp, qnrS1*, and *bla*CTX-M genes in transconjugants.
